# Homologous chromosomes in asexual rotifer *Adineta vaga* suggest automixis

**DOI:** 10.1101/2020.06.16.155473

**Authors:** Paul Simion, Jitendra Narayan, Antoine Houtain, Alessandro Derzelle, Lyam Baudry, Emilien Nicolas, Marie Cariou, Nadège Guiglielmoni, Djampa KL Kozlowski, Florence Rodriguez Gaudray, Matthieu Terwagne, Julie Virgo, Benjamin Noel, Patrick Wincker, Etienne GJ Danchin, Martial Marbouty, Bernard Hallet, Romain Koszul, Antoine Limasset, Jean-François Flot, Karine Van Doninck

## Abstract

The several hundreds of species of bdelloid rotifers are notorious because they represent an ancient clade comprising only asexual lineages^1^. Moreover, most bdelloid species have the ability to withstand complete desiccation and high doses of ionizing radiation, being able to repair their DNA after massive genome breakage^2^. To better understand the impact of long-term asexuality and DNA breakage on genome evolution, a telomere-to-tolemere reference genome assembly of a bdelloid species is critical^3, 4^. Here we present the first, high quality chromosome-scale genome assembly for the bdelloid *A. vaga* validated using three complementary assembly procedures combined with chromosome conformation capture (Hi-C) data. The different assemblies reveal the same genome architecture and using fluorescent *in situ* hybridization (FISH), we demonstrate that the *A. vaga* genome is composed of six pairs of homologous chromosomes, compatible with meiosis. Moreover, the synteny between homoeologous (or ohnologous) chromosomes is also preserved, confirming their paleotetraploidy. The diploid genome structure of *A. vaga* and the presence of very long homozygous tracts show that recombination between homologous chromosomes occurs in this ancient asexual scandal, either during DSB repair or during meiotic pairing. These homozygosity tracts are mainly observed towards the chromosome ends in the clonal *A. vaga* suggesting signatures of a parthenogenetic mode of reproduction equivalent to central fusion automixis, in which homologous chromosomes are not segregated during the meiotic division.

---

Bdelloid rotifers are a notorious clade of ancient asexual animals. However, both its longevity (>60 My) and diversity contradicts the expectation that obligatory parthenogenetic animal lineages are evolutionary dead-ends. Historical observations (or lack thereof) have produced a consensus that bdelloid rotifers are strictly parthenogenetic without any meiosis (e.g. no males or hermaphrodites^1^, ameiotic genome structure in the model species *Adineta vaga*^3^, apomictic oogenesis^5, 6^). However, recent studies brought doubt regarding the supposed absence of meiotic recombination in these microscopic animals. These include a drop of linkage disequilibrium with increasing distance between loci in *A. vaga*^7^, signatures of gene conversion^3, 8^, heterozygosity levels falling within the range observed for sexual metazoans^3, 4, 9^ and reports of allele sharing between bdelloid individuals^7, 10–12^.

In addition to its asexual evolution, the bdelloid rotifer *A. vaga* also became a model species for its extreme resistance to desiccation and radiation, with implications for space research. Both prolonged desiccation and radiation induce oxidative stress and massive genome breakage that *A. vaga* seem to handle well, maintaining high survival and fecundity rates while efficiently repairing its DNA double strand breaks (DSBs)^2^. The exact nature of their DSB repair mechanism remains unknown, but homologous recombination (HR) should be privileged in the proliferating cells in order to maintain genome stability and long-term survival.

Here we used state-of-the-art technologies and assembly methods, combining short reads, long reads and chromosome conformation capture (Hi-C) in *A. vaga*, to produce both haploid and phased chromosome-scale assemblies. We uncovered diploidy and homologous chromosomes in *A. vaga*, challenging our view on many fundamental aspects of their biology by reviving the possibility of meiosis and meiotic recombination in these ancient asexuals. Characterizing homologous chromosomes as potential templates for DNA repair through HR reconciles bdelloids genomics with their extreme desiccation resistance. This seminal high-quality genome for *A. vaga* complements those of *Caenorhabditis* and *Drosophila* species for comparative biology within protostomians.

## A diploid and paleotetraploid genome

Combining short and long sequencing reads (see Supplementary Table 1) with multiple independent genome assembly procedures relying on different assumptions regarding ploidy levels (Bwise^13^, Flye^14^ and Falcon^15^) followed by Hi-C scaffolding with instaGRAAL^16^, revealed similar chromosome-scale assemblies and comparable genome size estimations (Figure 1A). They all converged towards concordant genomes structures, as the six longest scaffolds from the haploid assembly (hereafter named AV19) were each colinear to exactly two long scaffolds from the phased assembly (Figure 1B). All pairwise alignments of the three independent assemblies (i.e. “haploid”, “diploid” and phased”, see Figure 1A) confirmed a chromosome-scale synteny (Supp. Fig. S1, S2, S3). In order to validate these assemblies, *in situ* hybridization analysis was performed with three pairs of fluorescent probes (FISH) targeting distinct halves of three chromosomes from the AV19 genome (Figure 1B, right side). The FISH showed that each probe pair (one green and one red) labelled the two same chromosomes with no or little overlap between both signals (Figure 1C). Chromosome painting was perfectly consistent with our chromosome-scale assemblies demonstrating that the *A. vaga* genome is diploid and composed of six pairs of colinear homologous chromosomes. Our new AV19 assembly was compared to the previous draft genome assembly, hereafter named “AV13”^3^. Upon investigation, none of the colinearity breakpoints and palindromes previously described were retrieved (Supp. Fig. S4, S5). This chromosome-scale colinearity was also observed between pairs of homoeologous (or ohnologous) chromosomes in the AV19 genome, a signature confirming the previously reported paleotetraploidy of *A. vaga* genome^3, 4, 8^ (in blue on Figure 1B). The three chromosome pairs 1, 2 and 3 are indeed homoeologous (or ohnologous) to the three pairs 4, 5 and 6, respectively. *A. vaga* is thus a diploid, paleotetraploid species in which synteny is well conserved along homoeologous chromosomes with 40% of the genes having retained their homoeologous copy (see Materials & Methods section).

**Figure 1.**
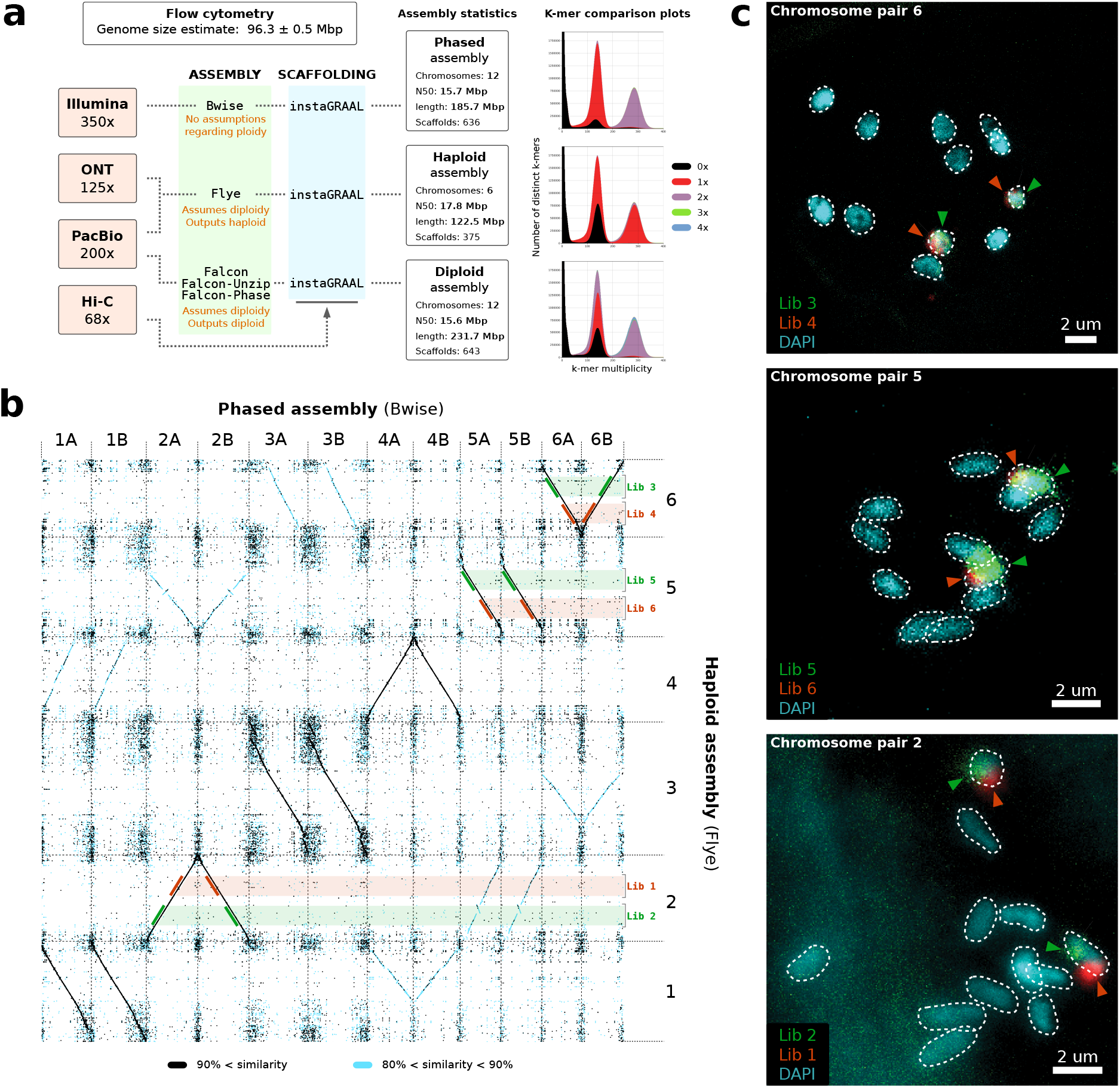
Genome structure of *Adineta vaga* is diploid. a) Outline of the three genome assembly approaches underlined by different assumptions on genome ploidy with median read coverage for all sequencing technologies indicated on the left and estimated with respect to the AV19 haploid genome assembly. The genome size estimate of *A. vaga* obtained by flow cytometry is given, as well as the summary statistics of the genome assemblies and the ploidy levels (KAT plots of k-mers distribution). Number of chromosomes corresponds to the number of scaffolds longer than 10 Mbp. b) Dotplot of the pairwise colinearity between the Flye and Bwise genome assemblies using minimap2 and D-genies. Black: *>* 90% identity; Light blue: *>* 80% identity. A schematic view of the design and position of three pairs of FISH libraries on colinear chromosome pair 6, 5 and 2. Karyotype of *A. vaga* with chromosome oligo painting (i.e. fluorescent *in situ* hybridisation) of three pairs of libraries designed on three chromosomes of an haploid assembly (designed as depicted in panel b).

## Homologous recombination in bdelloids

The discovery of homologous chromosomes in the oldest asexual animal clade known represents a major paradigm shift for studies of reproductive modes and led us to reconsider the possibility of homologous chromosome pairing in *A. vaga*. We measured and compared heterozygosity along the chromosomes of three *A. vaga* lineages cultured from a laboratory strain that never underwent desiccation or radiation and that were sequenced at three distinct timepoints (2009, 2015 and 2017, Figure 2A). Average SNP heterozygosity was around 1.5% (dark grey area on 2A, similar to previous studies^3, 4^). Importantly, we observed large homozygous regions (up to 4.5 Mb) in some isolates that were absent from others and seemed to accumulate with time (numbered tokens on Figure 2A). Given the genealogy of these laboratory populations, homozygous regions are the result of independent losses of heterozygosity (referred to as homozygotisation events, Figure 2B). These large-scale homozygosity tracts are absent from an *A. vaga* lineage sampled from the wild (using short-reads mapping, see Supp. Fig. S6) and might therefore be specific to laboratory culturing conditions, possibly through bottlenecks or reduction of selection leaving homozygotisation accumulation, usually considered deleterious, unchecked. Regardless of the underlying molecular mechanisms producing these homozygotisation events being mitotic, meiotic, or both, the existence of these events alone is a testament that molecular processes involving recombination of homologous chromosomes do occur in the germline of *A. vaga*.

**Figure 2.**
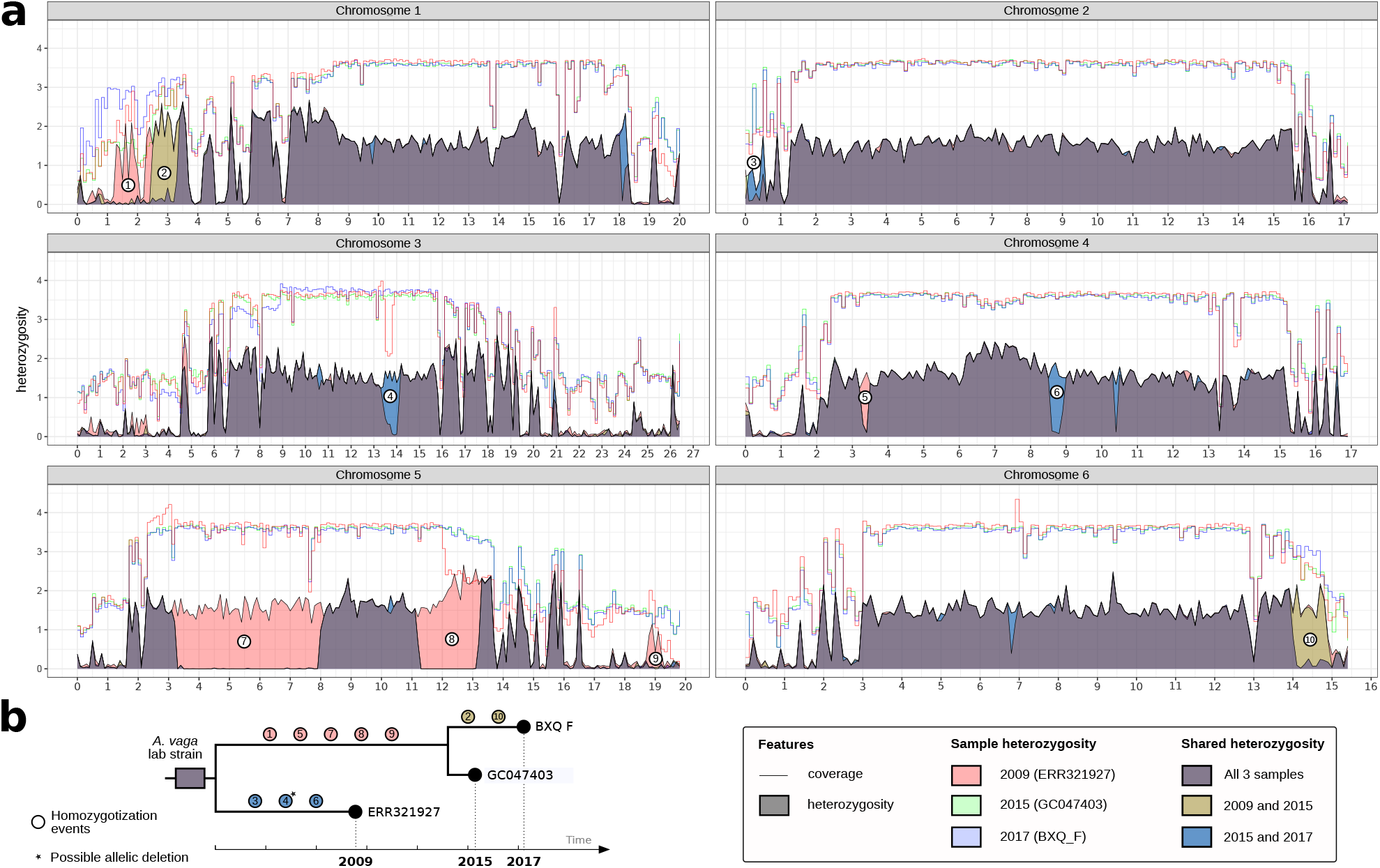
Dynamics of heterozygosity in *Adineta vaga*. a) Heterozygosity changes between three samples from the same strain along the six chromosomes. Lines indicate short read coverage and filled areas indicate heterozygosity. Superposition of filled areas indicate shared heterozygosity among samples and produces additional colors described in the legend. Sample ERR321927 correspond to short-read data used to assemble the first version of *A. vaga* genome (i.e. AV13^3^). Chromosome lengths are represented in Mb. b) Schematic evolutionary reconstruction of herozygosity changes among 3 samples from the same initial lab strain. Note that every sample had its own independent evolution, although its exact duration is unknown.

## Meiotic versus mitotic recombination

The presence of homologous chromosomes in *A. vaga* now means that bdelloid rotifers being ameiotic is only suggested by a mere absence of positive evidence for meiosis. Furthermore, reported genomic features are compatible with the occurrence of meiotic recombination: *A. vaga* genome is diploid (Figure 1) with an heterozygosity level in par with other metazoans^3, 4, 9^ (Figure 2A, Supp. Fig. S6), linkage disequilibrium decays with distance in *A. vaga*^7^, meiotic genes are present^3, 4, 17^, signatures of gene conversion have been reported^3, 8^ and here we show homozygotisation processes possibly reminiscent of crossing-overs (CO) (Figure 2). These homozygotisation events are both large-scale and frequently located at subtelomeric regions (i.e. events 1, 2, 3, 9 and 10 on Figure 2A), a signature expected in case of central fusion automixis. Under such form of parthenogenesis, maternal heterozygosity is retained because homologous chromosomes are not segregated during oogenesis (i.e. the reductional division is incomplete or the meiotic products of this first division get fused). In such situation, COs can produce homozygous chromosomal regions distal to the recombination event^18, 19^.

Alternatively, HR between homologous chromosomes producing long homozygotisation events (Figure 2) may also occur during mitotic DSB repair^20^. Most bdelloids are known for their capacity to endure and repair extensive genome breakage, induced by prolonged desiccation in their semi-terrestrial habitats or by high doses of ionizing radiation^2^. DSBs can be lethal and drive genetic instability if not repaired. While DSB repair mechanisms requiring no homology, such as non-homologous end-joining (NHEJ), could still potentially be used in the somatic cells of eutelic bdelloid rotifers^21^, it is likely that the germline is maintained by HR repair pathways, using the sister chromatid or homologous chromosome as template. These would ensure that the oocytes pass the molecular checkpoints of DNA integrity during oogenesis^22^ and that the genome structure is maintained even after experiencing massive DSBs. Intriguingly, an analysis of gene enrichment in the two large homozygous tracts observed on chromosome 5 (see Figure 2A, samples from 2015 and 2017) showed a highly significant enrichment of DNA repair function compared to the rest of the genome (see Supplementary Table 2). It is still unclear whether this enrichment is due to evolutionary processes or to chance, as every chromosome presents some enriched function when compared to the five other chromosomes (see Supplementary Table 2). Several homology-driven repair mechanisms exist that could potentially be at play to produce homozygosity in bdelloid rotifers, and their determination should be the focus of future studies. Overall, these mechanisms could be invoked in both a meiotic and/or a mitotic context and the general question whether reproductive mode, desiccation resistance mechanisms, or both, mainly shaped the genome of bdelloid rotifers remains open.

## Automixis is not scandalous

The bdelloid rotifer clade, having diversified into more than 400 asexual morphospecies, was deemed scandalous because it was interpreted in a context of apomixis, devoid of both meiosis and recombination^23^. Under our new hypothesis of plausible automictic parthenogenesis with modified meiosis, the scandalous longevity of the bdelloid clade is solved by the existence of within-individual recombination. Indeed, within-individual recombination partially circumvents the expected deleterious consequences of strict clonality^3, 24^ (i.e. Muller’s ratchet) by decoupling deleterious mutations from beneficial ones, improving the efficiency of selection^25, 26^. It has indeed already been argued that automictic lineages could be “superior to both clonal and outbreeding sexual populations in the way they respond to beneficial and deleterious mutations”^18^. Modelisation of automictic population genetics indicates automixis is theoretically compatible with long-term evolutionary success, as it can lead to a higher neutral genetic diversity than in sexual lineages, a lower mutational load than in both clonal and sexual lineages and a lower genetic load than in sexuals subjected to overdominant selection^18^. We argue here that it is likely that parthenogenesis in *A. vaga* relies on a modified meiosis genetically equivalent to central fusion automixis as in several other metazoans (*Aspidoscelis uniparens*^27^, *Apis mellifera capensis*^28^, *Daphnia pulex*^29^, *Daphnia magna*^30^ or *Artemia parthenogenetica*^31^. If our hypothesis holds true, then bdelloid rotifers might switch status from scandalous ancient asexuals (referring here to apomixis) to successful ancient automictic.

## High HGT content

The dynamics of foreign DNA has been hypothesized to potentially play an important role in bdelloid evolution, notably because genetic transfers could be a way to circumvent some of the deleterious effects of obligate apomixis through the integration of non-metazoan DNA, thereby possibly triggering adaptation^2, 4, 32^. We developed an innovative HGT detection tool, Alienomics, which combines many criteria such as gene taxonomy, GC content, coverage and synteny to accurately detect HGTs (see M&M). Using Alienomics on AV19 genome, we found 2,615 non-metazoan gene transfers (8.9% of all genes) in *A. vaga*, confirming previous reports of highest HGT content within metazoans^3, 4, 33, 34^. We found that HGTs are not restricted to subtelomeric regions, as previously reported^33^, but somewhat homogeneously distributed along chromosomes with local hotspots (Figure 3A). Surprisingly, two HGT hotspots were found in homoeologous chromosomes (Figure 3B), suggesting that this HGT hotspot predates the ancient tetraploidization of bdelloids^8, 34^. Further analyses will be necessary in order to determine whether the horizontally acquired genes in these regions predate the origin of bdelloids or if only the HGT hotspot itself, but not its content, has been conserved across homoeologous chromosomes. A previous study found that about 80% of the HGTs across 4 species including *A. vaga* were co-orthologs, suggesting an ancient, shared origin of these foreign genes in bdelloid rotifers, at least as old as the divergence between *Adineta* and *Rotaria* species^4^. Finally, we report an enrichment of two functions important for desiccation resistance when comparing HGT genes versus the entire genome: carbohydrate metabolic processes and oxidation-reduction processes (see Supplementary Table 2), thereby confirming previous findings^2, 3, 32^. While HGTs might not be needed to explain the longevity of asexual rotifers, our results reinforce the idea of an evolutionary importance of HGTs in bdelloids, possibly predating bdelloid origin and thus potentially linked to the acquisition of obligatory parthenogenesis and/or desiccation resistance.

**Figure 3.**
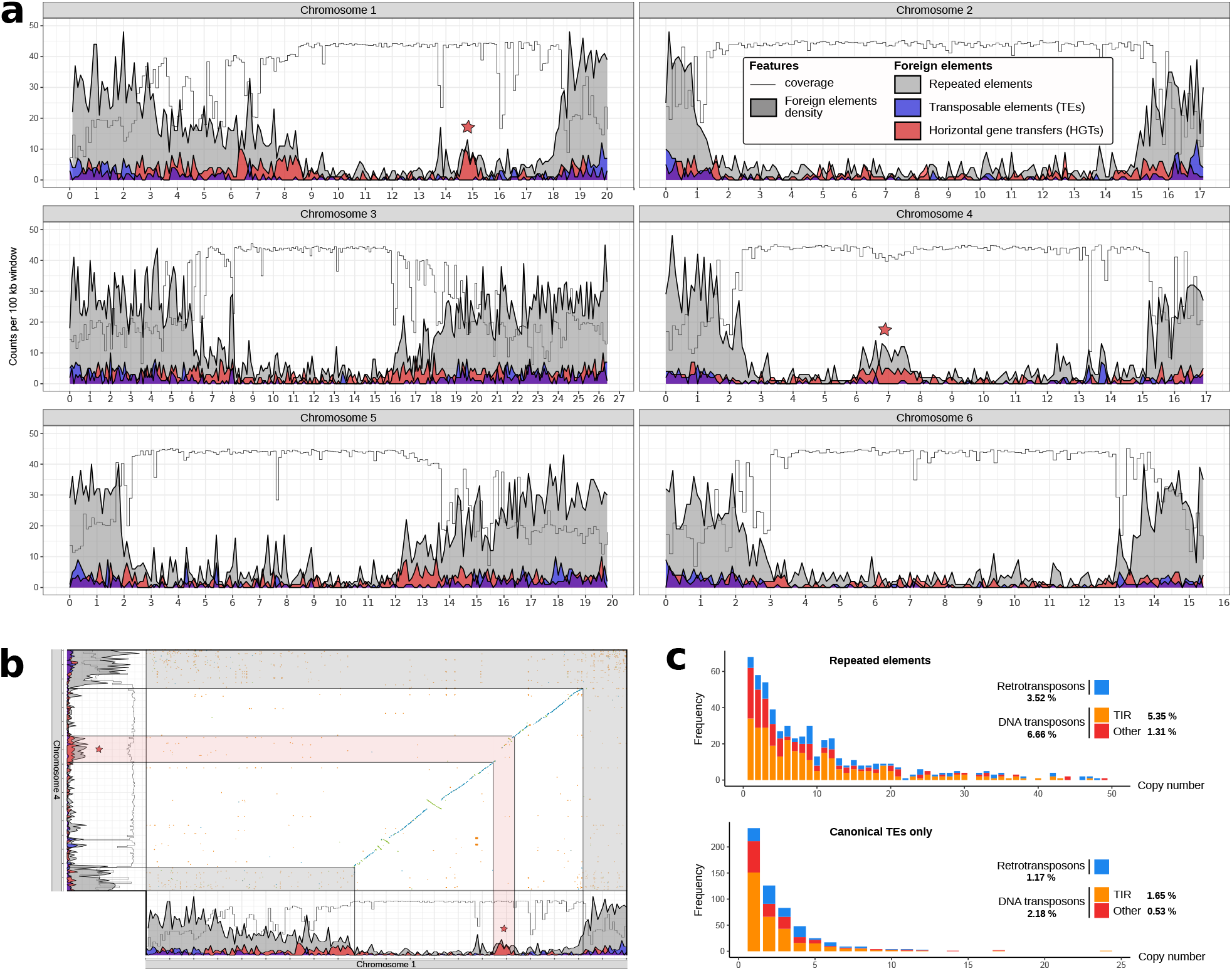
Quantity and distribution of foreign DNA along chromosomes. a) Density of HGTs, repeated elements and TEs along the 6 chromosomes of the AV19 genome assembly (x-axis in Mb). Red stars indicate hotspots of HGT and repeated sequences. b) Ancient conservation of HGT hotspot localization, highlighted in red, between homoeologous chromosomes 1 and 4. Non conserved subtelomeric regions are highlighted in grey. c) Composition and number of copies of repeated elements (top) and canonical TEs (bottom)

## Repeated and transposable content

Two approaches were combined to annotate both repeated elements and canonical TEs (i.e. EDTA and REPET pipelines, see Supp M&M). Repeated elements and canonical TEs respectively covered about 10.7% and 3.3% of *A. vaga* genome (in line with the ca. 3% TEs previously reported for this species^3, 4^) and most consensus sequences were found in low copy numbers (6*x*) as previously described^3, 4, 35^ (Figure 3C). Tandem inverted repeats (TIRs) DNA transposons (e.g. Class-II) were quantitatively dominant among the low amount of transposable elements in *A. vaga* genome (i.e. 53% of all repeats), in line with previous findings^3, 35^ (Figure 3C).

## Dynamics of foreign DNA

Past studies repeatedly suggested a preferential co-localisation of HGTs and TEs^33, 36, 37^ at subtelomeric regions in *A. vaga*^33, 35^. While repeated elements were found mostly at subtelomeric regions, canonical TEs were found both at the subtelomeres and along entire chromosomes (Figure 3A). We also found a weak, although significant, correlation of co-localization between repeated elements and HGTs when looking at genomic windows of 20kb or 100kb (Pearson’s correlation factor of 0.15 and 0.36, respectively, both with pvalues of 2.2*e*^−^16). A co-localization between HGTs and repeated elements can be observed at the ancient HGT hotspot well-within chromosomes (Figure 3A,B). Of note, the coverage of Illumina reads in the wild *A. vaga* lineage drops at these two hotspots S6. This suggests that while the localization of these hotspots of foreign DNA is ancient, part of their content (repeated elements, HGTs, or both) diverged more recently between the two *A. vaga* lineages. Given the strong association between repeated elements and telomeres in *A. vaga*, it is tempting to interpret the hotspots of repeated elements and HGTs as signatures of past chromosomal fusions, the hotspots representing ancient telomeres. Testing this hypothesis will however require additional chromosome-scale genome assemblies from other bdelloid species.

## Allele sharing in bdelloids

Lastly, whether bdelloid rotifers exchange genetic content and whether this is done through sexual reproduction or horizontal transfers (during desiccation) is still debated^7, 10–12, 38, 39^. Since we demonstrated that *A. vaga* genome structure is compatible with a meiotic parthenogenetic reproductive mode, there are now three main hypotheses that could explain the recurrent patterns of allele sharing recently reported: i) allele sharing is due to undetected contamination between cultures in all recent studies, either during colony culture itself or during sample preparation for sequencing^40–44^; ii) allele sharing results from facultative sex (i.e. cyclical parthenogenesis), with sex events being cryptic enough so that males, sperm or fertilization were never observed, but frequent and abundant enough so that every sampled population studied was sufficient to detect it^10, 12^; iii) allele sharing is the result of horizontal genetic transfers between bdelloid individuals through unknown molecular mechanisms, possibly associated to desiccation^2, 11, 38, 45, 46^ and potentially enabling integration of non-metazoan genes. While allele-sharing patterns remains highly debated, these three hypotheses are all compatible with automictic parthenogenesis in *A. vaga* and an ambitious sampling and population genomics investigation of wild *A. vaga* is ongoing to try to provide answers.

## Limitation of the current reference assembly

Pre-print readers will have noticed some aspects still potentially problematic with the AV19 genome assembly. First, its genome size estimate is larger than the robust estimation obtained by flow cytometry (Figure 1A). Second, the coverage distribution across all sequencing technologies diminishes to ~ 50% of the expected coverage at telomeric regions (Figure 2). After further investigations, it appears that both these issues are the consequence of remaining uncollapsed haplotypes in the AV19 haploid assembly^47^. We recently solved this issue and already produced an improved haploid genome assembly (AV20) that we are now analyzing. While we are confident that the results of this pre-print will not change, or only subtly, an upgraded version of this article is under way.

## Conclusion

This high-quality chromosome-scale assembly, still lacking in many model organisms, firmly establishes *A. vaga* as a ideal system to study long-term parthenogenetic evolution and recovery from genome breakage. Homologous chromosomes are present in bdelloids and heterozygosity changes across samples are only compatible with the existence of long-range homologous recombination, suggestive of homologous chromosome pairing. Our work reinforces the hypothesis that recombination is critical for lineage longevity and that ancient asexual animals without any form of meiosis or recombination might not exist at all. Meiotic recombination could be key both for faithful DSB repair after desiccation and for circumventing the deleterious effects often associated with asexuality. The *Adineta vaga* species is of prime interest for evolutionary biology and this genome will be a precious tool to unravel its population biology as well as its extreme resistance to desiccation and ionizing radiation.

## Materials and methods

### Strain culture, Library preparation and DNA sequencing

We continuously cultivated *A. vaga* individuals from AD008 strain (i.e. same strain as in^3^, COI sequence accession number is KM043184) since 2007 in Petri dishes using Spa water, feeding them with sterile extract of lettuce juice and stocking well-grown cultures at −80°C. *A. vaga* individuals were thawed before proceeding to DNA extraction using QIAGEN Gentra Puregene Tissue Kit. Genomics Core (UZLeuven) produced PCR-free 250-bp paired-end Illumina reads that were sequenced with a coverage of approximately 350x on a HiSeq 2500 sequencing platform. The same procedure was followed in order to obtain high molecular weight DNA using Macherey-Nagel NucleoBond HMW procedure that was subsequently sent to the Genomics Core (UZLeuven) to generate a coverage of 200x of PacBio RSII sequencing data. Around 30 μg of high molecular weight DNA was also extracted from living *A. vaga* individuals using the QIAGEN Gentra Puregene Tissue Kit and then sent to the Genoscope sequencing center (François Jacob Institute of Biology) which produced 5 ONT libraries, each starting from 2 to 5 μg of DNA, using the 1D ligation sequencing kit (SQ-LSK108) and R9.4 (or R9.4.1) flowcells. This resulted in a coverage of 125x long-reads using Oxford Nanopore Technology (ONT). Samples ID and SRA accession numbers are detailed in Supplementary Table S1.

### Chromosome conformation capture

The Hi-C library construction protocol was adapted from^48, 49^. Briefly, individuals from the *A. vaga* AD008 strain were chemically cross-linked for 20 min at room temperature and 30 min at 4°C (with gentle stirring) using formaldehyde (final concentration: 5% in milliQ water; final volume: 50 ml). After fixation the sample is centrifuged for 10 min at 4000 rpm at 4°C. The formaldehyde was then quenched for 5 min at RT and 15 min at 4°C (with gentle stirring) by adding 50 ml of 250mM glycine. The cells were recovered by centrifugation for 10 min at 4000rpm at 4°C, supernatant is removed and pellet stored at −80°C until use. The Hi-C library was then prepared as follow. Cells were resuspended in 1.2 mL of 1X DpnII buffer (NEB), transferred to a VK05 tubes (Precellys) and disrupted using the Precellys apparatus and the following program ([20 sec – 6000 rpm, 30 sec – pause] 9x cycles). The lysate was recovered (around 1.2 mL) and transferred to two 1.5 mL tubes. SDS was added to a final concentration of 0.3% and the 2 reactions were incubated at 65°C for 20 minutes followed by an incubation of 30 minutes at 37°C. A volume of 50 μL of 20% triton-X100 was added to each tube and incubation was continued for 30 minutes. DpnII restriction enzyme (150 units) was added to each tube and the reactions were incubated overnight at 37°C. Next morning, reactions were centrifuged at 16,000 x g for 20 minutes. The supernatants were discarded and the pellets were resuspended in 200 μL of NE2 1X buffer and pooled (final volume = 400 μL). DNA extremities were labelled with biotin using the following mix (50 μL NE2 10X buffer, 37.5 μL 0.4 mM dCTP-14-biotin, 4.5 μL 10mM dATP-dGTP-dTTP mix, 10 μL Klenow 5 U/μL) and an incubation of 45 minutes at 37°C. The labelling reaction was then split in two for the ligation reaction (ligation buffer – 1.6 mL, ATP 100 mM – 160 μL, BSA 10 mg/mL – 160 μL, ligase 5 U/μL – 50 μL, H2O – 13.8 mL). The ligation reactions were incubated for 4 hours at 16°C. After addition of 200 μL of 10% SDS, 200 μL of 500 mM EDTA and 200 μL of proteinase K 20 mg/mL, the tubes were incubated overnight at 65°C. DNA was then extracted, purified and processed for sequencing as previously described^48^. Hi-C libraries were sequenced on a NextSeq 550 sequencer (2×75 bp, paired-end Illumina NextSeq with the first ten bases acting as barcodes^50^).

### Bwise assembly

The Bwise assembler v0.1 (https://github.com/Malfoy/BWISE) was used on high-coverage Illumina data (sample ID GC047403, see Supplementary Table 1) to produce a draft phased genome assembly. We selected a Kmer size parameter of 63 (-k 63) as this produced the most contiguous assembly over the range of tested Kmer sizes: 63, 73, 101, 201. Other parameters were left as default. Bwise rests on a different paradigm than most assemblers: it starts by generating a de Bruijn graph from the reads to assemble^51^, then cleans the graph by removing tips caused by sequencing errors^52^, remaps the initial reads on this corrected de Bruijn graph^53^, transforming them in super-reads^54^. Finally, the resulting super-reads are assembled in a greedy fashion whenever they overlap unambiguously by one or several unitigs. This approach was devised in order to produce an assembly that reflects faithfully the unknown ploidy level of the organism sequenced. Therefore, Bwise will produce haploid assemblies whenever the organism sequenced is haploid, diploid assemblies whenever the organism sequenced is diploid, triploid… etc.

### FLYE assembly

The Flye genome assembly software version 2.5^14^ was run using both PacBio (350 X coverage) and ONT (125 X coverage) long reads, with default settings (see Supplementary Table 1). Flye is a OLC (overlap-layout-consensus) long-read assembler especially designed to handle repeats. During the layout step, it uses a repeat graph allowing for approximate sequence matches, therefore tolerating long reads errors. Briefly, at the beginning of the layout step the initial disjoint, error-prone contigs (disjointigs) are concatenated in a random order. An assembly graph (the repeat graph) is subsequently constructed from the repeat plot of the concatenates and each edge of the graph is classified according to its multiplicity (unique or repetitive). Mapping the reads on that repeat graph allows resolving bridged repeats and subsequently unbridged repeats. Flye solves repeats not fully covered by reads by identifying variations between copies and matching each read to its respective copy. Flye is, however, not able to solve unbridged repeats if no variation between copies exists.

### FALCON assembly

The *de novo* assembly of *Adineta vaga* genome was carried out with diploid-aware long-read assembler FALCON version 0.7.0, FALCON-Unzip and partial FALCON-Phase (only FALCON-Phase Workflow steps 1, 2 and 3)^15^. Prior to the assembly, Canu error correction module was used for read error correction based on raw PacBio reads. The FALCON software is highly optimised for eukaryote genomes, and uses hierarchical genome assembly process (HGAP). More specifically, reads longer than 15 kb were selected by Falcon as “seed” reads to generate consensus sequences with high accuracy. The pre-assembly steps in FALCON uses DALigner to do all-by-all alignments of the corrected PacBio reads. Long reads were then trimmed at regions of low coverage with FALCON sense parameters (-minidt 0.70 -mincov 4-maxnread 200) and sensitive DALigner parameters were selected (-h60 −e.96 −l500 −s1000) for pre-assembly process. The FALCON pre-assembly resulted in 331 primary contigs of total length 125 Mb, contig N50 of 6 Mb and an additional 36 Mb of “associate contigs” that represent divergent haplotypes in the genome. FALCON-unzip was then used to phase the pre-assembly, producing contiguous leading contigs (named “primary”) and associated contigs (i.e. phased, alternate haplotypes). The genome assembly was polished as part of the FALCON-Unzip pipeline using haplotype-phased reads. The haplotigs contain one of the two allelic copies of the heterozygous regions; in this respect, the haplotigs serve as phasing information for the haploid representation. The FALCON-Unzip assembly had 241 primary contigs and 999 haplotigs. FALCON-Phase (https://github.com/phasegenomics/FALCON-Phase) was developed to resolve haplotype switching in diploid genome assemblies. The FALCON-Phase haplotig placement defines phased blocks in the FALCON-Unzip assembly. The Falcon-Phase Workflow steps 1 and 2 were used to place the haplotigs along primary contigs. Once the haplotig placement file and phase block pairings are done, the primary contigs are cut up into very small pieces at phase block boundaries with Falcon-phase workflow step 3.

### Scaffolding with instaGRAAL

Hi-C contact maps were generated from paired-end reads using the hicstuff pipeline for processing generic 3C data, available at https://github.com/koszullab/hicstuff. The backend uses the bowtie2 aligner run in paired-end mode (with the following options: -{}-maxins 5 -{}-very-sensitive-local). We discarded alignments with mapping quality lower than 30. The remainder was converted to a sparse matrix representing contacts between each pair of DpnII restriction fragments. The instaGRAAL program^16^ was used in conjunction with the contact maps to scaffold the genomes. Prior to running it, restriction fragments are filtered based on their size and total coverage. Fragments shorter than fifty base pairs are discarded. Then, fragments whose coverage lesser than one standard deviation below the mean of the global coverage distribution are also removed from the initial contact map. These fragments were reintegrated later after the scaffolding step. The instaGRAAL scaffolder uses a Markov Chain Monte Carlo (MCMC) method: briefly, the contact data is fitted on a simple three-parameter polymer model. The 3D contacts are exploited and used by the program to infer the relative 1D positions of the sequences and thus the genome structure. To do so, the program attempts to perform a number of operations between each sequence and one of its neighbours (*e.g.* flipping, swapping, merging or splitting contigs) and the operation is either accepted or rejected with a certain probability depending on the likelihood shift. The model parameters are then also updated and a new iteration begins. A set of computations whereby every sequence of the genome has been iterated over this way is called a *cycle*. The scaffolder was run for 100 cycles on each genome, after which convergence in both genome structure and model parameters was evidently apparent. The scaffolded assemblies were then refined using instaGRAAL’s instaPolish module, with the aim of correcting the small artefactual inversions sometimes produced by instaGRAAL.

### Post-treatment of scaffolded assemblies

Diploid assembly post-treatment (using FALCON): we used the repeat-aware finisherSC tool^55^ to upgrade the *de novo* phased genome assembly of *Adineta vaga*. Final round of polishing were performed with the Pilon corrector using Illumina data (sample ID GC047403, see Supplementary Table 1). Phased assembly post-treatment (using Bwise): to resolve a remaining fragmentation of one single chromosome (i.e. chromosome 5B) after scaffolding with instaGRAAL based on Hi-C data, we established a novel comparative approach that incorporates computational methods to transform fragmented contigs into near-chromosome fragments. First, Bwise contigs were aligned against themselves using NUCmer v4.0^56^. Ploidy pairing was evaluated using the online visualization tool, DOT (https:/dnanexus.github.io/dot/) and we were able to anchor fragmented contigs into a single chromosome using its homologous template (i.e. chromosome 5A).

### Ploidy, synteny and colinearity among the three *A. vaga* genome assemblies

Genome assembly tools rely on various assumptions including the ploidy level of the organism under study. In order to circumvent potential impact of such ploidy assumptions on genome structure, we compared our three new genome assemblies. First we evaluated the classical genome assembly statistics using in-house script (see Figure 1A). We then used the illumina reads (i.e. GC047403, see Supplementary Table 1, as input for the *comp* function of the KAT software^57^ which uses k-mers distribution in order to explore ploidy levels of *A. vaga* genomes (see 1A). Genomes were aligned pairwise using nucmer 3.1 (using –maxmatch option)^58^, the results of which were converted into paf format using minimap2 paftools script^59^. We then used D-GENIES as a stand-alone^60^, modifying identity percentage thresholds to 1.0/0.9/0.8/0.5 to visualise the three pairwise alignment as dotplots (see Figure 1 and Supp. Fig. S1, S2 and S3).

### Genome size estimation

The genome assemblies produced by all three methods (Bwise, Flye and Falcon) were markedly smaller than expected based on the generally admitted genome size of 0.25 pg per (non-reduced) oocyte (http://www.genomesize.com/result_species.php?id=5369), equivalent to 244 Mbp for a diploid assembly or 122 Mbp for a haploid assembly. As there is considerable confusion in the literature considering the genome size of *Adineta vaga* (*e.g.* report of a nuclear DNA content of about 0.7 pg^61^, nearly 3 times higher than in the Animal Genome Size database although the entry there refers to this article), we decide to perform an independent assessment of the genome size of *Adineta vaga* using flow cytometry, with *Arabidopsis thaliana* ecotype Colombia (for which a genome size of 157 Mbp was previously measured^62^ as a genome-size standard for comparison. Nuclei from both species were isolated according to the protocol from the Cystain Pi absolute T (SYSMEX #05-5023) kit. Briefly, we chopped them together in the same extraction buffer (500 *μ*L), after which the material was filtered through a 30 *μ*m nylon membrane. After RNAse treatment (80 *μ*g/ml), the DNA was labeled for 1h in the dark with 2 ml of staining buffer containing 120 *μ*L of propidium iodide. The labeled nuclei were then analyzed on the CyFlow Space flow cytometer (Sysmex) of the research unit “Evolutionary Biology & Ecology” of the Université libre de Bruxelles (ULB). We used a blue laser with an excitation wavelength of 488 nm. The whole procedure was performed three times on different days, using different batches of rotifers and leaves from a different *A. thaliana* plants every time, and the. FCS files were analyzed using the FlowJo v10.6.2 software. The estimated haploid genome size is presented in Supp. Fig. S7.

### Chromosome painting (FISH)

To assess the colinearity between two chromosomal markers, FISH experiments were performed on samples containing well resolved condensed chromosomes. As bdelloids are eutelic, such condensed chromosomes are only found in embryos undergoing nuclear divisions. Particularly, young embryos containing only few nuclei usually exhibit the nicest karyotypes^63^. To collect young, ideally one-cell, embryos, about 200 rotifers bearing a single egg were first isolated in a petri dish containing a 1% agarose pad and ice-cold Spa® spring water. The agarose pad avoids the embryos to stick at the bottom of the plate and ease their isolation. The rotifers were starved for 24 hours at 4°C and, the next day, about half of the water was removed and replaced by the same volume of fresh water at RT containing lettuce filtrate. Rotifers were incubated at 25°C and, about 3 hours later, all individuals were laying eggs almost synchronously. Immediately after laying, the eggs were collected and fixed in methanol (Merck Millipore®, 1070182511): acetic acid glacial (VWR™, 20104-243) (3:1) solution on ice. After isolation of all eggs, they were collected by centrifugation (14,000 rpm, 2 min, RT), fixed again with methanol: acetic acid glacial (3:1) and stored at 4°C until slide preparation. About 100 embryos bearing one or few nuclei can be collected by this method. For the FISH probe synthesis, we used the Oligopaint strategy that consists in the use of libraries of short single-stranded oligonucleotides (oligos) that are fluorescently labeled to visualize megabases (Mbs) of genomic regions^64^. The design of the probes was performed using the OligoMiner pipeline^65^ that selects for oligos having similar parameters such as melting temperature (Tm) or the absence of secondary structures. The selected oligos have a 30-42 nt region of genomic homology with a Tm of 42°C flanked by constant nongenomic sequences at the 5’ end (5’-ccc-gcg-tta-acc-ata-cac-cg-3’) and at 3’ end (5’-ggt-agc-cac-acg-ctt-cga-tg-3’). These sequences are necessary for the labeling and the amplification of the libraries by PCR (see below). We ordered 6 libraries from GenScript®: (i) library 1 (7.7k oligos) targets the chromosomes 2a/b from 2 to 6 Mbs; (ii) library 2 (9.2k oligos) targets the chromosomes 2a/b from 13 to 16 Mbs; (iii) library 3 (7.7k oligos) targets the chromosomes 6a/b from 2 to 6 Mbs; (iv) library 4 (8.0k oligos) targets the chromosomes 6a/b from 8 to 12 Mbs; (v) library 5 (7.9k oligos) targets the chromosomes 5a/b from 3 to 7 Mbs; and (vi) library 6 (7.8k oligos) targets the chromosomes 5a/b from 9 to 13 Mbs. The probes were labeled and amplified according to the ‘One-day’ probe synthesis protocol using lambda exonuclease described in^66^ (https://oligopaints.hms.harvard.edu/protocols). The oligo libraries were first amplified and labeled by PCR. Twenty-four PCR reactions (24 x 50 *μ*l) were performed with 1 U of Q5 high-fidelity polymerase (New England Biolabs®, M0491), 200 *μ*M dNTPs, 0.5 *μ*M of fluorescently labeled forward primer (5’-/Fluo/ccc-gcg-tta-acc-ata-cac-cg-3’), 0.5 *μ*M of phosphorylated reverse primer (5’-/Phos/cat-cga-agc-gtg-tgg-cta-cc-3’), and 1.25 ng of Oligopaint library. The primers were ordered from IDT®. To perform the two-color FISH experiments, libraries 2, 3 and 5 were labeled with 5Atto488N (green) and the libraries 1, 4, and 6 were labeled with Atto565N (red). The PCR reactions were incubated at 98°C for 5 min, followed by 40 cycles of 30 sec at 98°C, 30 sec at 56°C, and 15 sec at 72°C, and a final extension at 72°C for 5 min. The PCR reactions were then collected and concentrated using the Zymo DNA clean concentrator kit (Zymo research®, D4032). The concentration was performed according to the manufacturer protocol and the libraries were eluted in 2,800 *μ*l of RNase/DNase free water. Lambda exonuclease (New England Biolabs®, M0262) was then used to hydrolyze the 5’-phosphorylated strand of the double-stranded amplicons. DNA eluant (2,200 *μ*l) was processed by 250 U of lambda exonuclease at 37°C for 30 min, and then stopped by incubation at 75°C for 10 min. The single-stranded labeled probes were finally cleaned up using the Monarch PCR & DNA cleanup kit (New England Biolabs®, T1030) following the oligonucleotide cleanup protocol. Probes were eluted in 20 *μ*l of RNase/DNase free water and stored protected from light at −20°C until use. The hybridization of the probes on embryos was adapted from previous protocols^63, 67^. At least 100 embryos stored in methanol: acetic acid glacial were dropped onto an uncoated and clean microscope slide (VWR™, 631-1550) and let dry on a wet paper for 30 min. Then, a cover slip (VWR™, 631-1572) was placed over the embryos and they were squashed by gentle pressure on the slide. All following treatments of embryos on slides were conducted in Coplin jars. Embryos were permeabilized in 0.1% saponin (Sigma-Aldrich®, 47036)/0.1% triton X-100 (Sigma-Aldrich®, T8787) in PBS (Lonza®, 17516Q) for 10 min, followed by 2 washes of 5 min in PBS. Samples were incubated for 20 min in PBS containing 20% of glycerol (Carl Roth®, 7530.1) and washed again 2 times in PBS. Slides were incubated for 5 min in 2x SSC (SSC 20X, Invitrogen 15557-036) supplemented with 0.1% of Tween-20 (Sigma-Aldrich®, P1379) (i.e., 2x SSCT), and then for 5 min in 2x SSCT supplemented with 50% of formamide (Sigma-Aldrich®, 47671). The slides were then put on top of a thermoblock at 92°C for 2.5 min and transferred in a Coplin jar containing 2x SSCT-50% formamide at 60°C for 20 min. The jar was then removed from 60°C and placed at RT for 1 hour. The hybridization mixture (50 *μ*l) composed of 2x SSC, 50% formamide, 1 ul of RNAse A (Sigma-Aldrich®, R4642), 10% dextran sulfate (Sigma-Aldrich®, S4030), and 10 *μ*l of each labeled oligo libraries, was placed on a clean cover slip and the slide containing the embryos was inverted onto this cocktail of hybridization. For the two-colors FISH, the oligo library 1 (red) was mixed with the oligo library 2 (green), the oligo library 3 (green) was mixed with the oligo library 4 red), and the oligo library 5 (green) was mixed with the oligo library 6 (red). The cover slip was sealed with rubber cement and let dry for 5 min at RT. The mounted slide was denatured at 92°C for 2.5 min on a thermoblock, transferred to a dark humidified chamber, and incubated O/N at 37°C. The next day, the cover slip was removed carefully from the slides. The slides were then washed in 2x SSCT at 60°C for 15 min, and in 2x SSCT at RT for 10 min. Chromosomes were counterstained for 20 minutes with 1 *μ*g/ml DAPI (4’,6-diamidino-2-phenylindole; ThermoFisher Scientific, D3571) in 2x SSC. Slides were washed twice in 2× SSC for 10 min, and mounted under a 24 × 32 mm cover slip in Mowiol 40-88 (Sigma-Aldrich®, 324590). Chromosomes and FISH signals were observed under a Leica TCS SP5 fluorescence confocal microscope using the 488 nm laser to capture the green signal, the 561 nm laser for the orange signal and the 405 nm laser line for the DAPI signal. Images were captured in Z-stacks with the LAS AF software and they were finally processed and analyzed with Fiji (ImageJ, version 2.0.0).

### Contamination check with blobtools

Contaminant reads from other organisms pollute the assemblies. Blobtools^68^ taxonomic partitioning was applied on the AV19 genome. Illumina paired-end HiSeq2500 HT reads were mapped on the assembly with bwa mem v0.7.17-r1188 (with the -Y parameters to mark supplementary alignments as soft-clipping)^69^ and bam files generated with samtools v1.9^70^. The taxonomic annotation of contigs was done with diamond blastx^71^ using the NCBI non-redundant protein database (nr) (downloaded the 11/23/18) and using bitscores instead of e-values. The blobplots (see Supp. Fig. S8) were obtained with blobtools v1.0 using default parameters. Note that no scaffolds were removed after this step, as the genome of *A. vaga* is expected to have a high content of true HGTs.

### Detecting homoeologous genes

The number of genes retaining their homoeologous counterpart was estimated by a similarity search of *A. vaga* proteome against itself using blastp^72^ (parameters as follows: -qcov_hsp_perc 30 -max_target_seqs 4) and by filtering out hits if their length was < 100 and if their idendity percentage was < 40, thus only retaining strong proteic alignments. These pairs of proteins were then filtered to only retained those for which the two proteins were respectively located onto two homoeologous chromosomes using in-house script. This procedure retained 12,976 genes out of the 32,378 proteins (i.e. 40.1%).

### Gene annotation

Gene prediction and annotation of AV19 genome were done according to current integrative approaches based on several independent lines of evidence. We first discarded scaffolds shorter than 1000 bp using funannotate clean function^73^. Repeats in the genome were then soft-masked using RepeatModeler^74^ and RepeatMasker^75^. RNA-Seq data from several cultured clones (see Supplementary Table 1) were used to produce *de novo* a transcriptomic assembly with trimmomatic^76^ and trinity^77^ both under default parameters. This transcriptomic assembly as well as additional RNA-Seq data directly mapping on the genome (see Supplementary Table 2) were used as input for the funannotate train function that wrap the PASA pipeline^78^ which relies on RNA-Seq to produce high quality annotations. Then, we used a combination of PASA annotations, *de novo* assembled transcriptome, metazoan BUSCO database and the proteic Uniprot database within funannotate predict function. This first produced *ab initio* predictions using Genemark-ES^79^, which were then used along with transcripts and proteic data to train Augustus to generate a second set of annotations. Lastly, it used Evidence Modeler as a weighted approach to combine annotations from PASA, Genemark and high quality predictions from Augustus into an integrated gene annotation set. We then used InterProScan5 in order to produce functional annotations to the predicted genes^80^ which were then used in combination with busco metazoan database using the funannotate annotate function with default parameters.

### Detecting HGTs with a new tool: Alienomics

We used a newly developed tool, named Alienomics, in order to detect putative Horizontal gene transfers (HGTs). This tool is being submitted and described in details elsewhere. Briefly, its approach first integrates several lines of quantitative evidence into a score for every predicted gene. This gene score is based on several blast results (i.e. against Uniref50 database, a user-defined set of closely-related reference genomes, bacterial rRNA database, BUSCO database) as well as on read coverage and GC content. It represents how “alien” or “self” a given gene is. We then use qualitative synteny information in order to discriminate if alien genes stemmed from contaminant or from HGT. For this, scaffolds are being given a score based on the integration of all the gene scores. This scaffold score represents whether it originated from a contaminant or from the genome under study. Synteny is then taken into account by comparing gene scores to their respective scaffold scores to validate a HGT. For example, an “alien” gene on a “self” scaffold corresponds to a HGT while an “alien” gene on an “alien” scaffold is a contaminant. Alienomics will soon be freely available for download and use on Github.

Within Alienomics, results for each criteria (e.g. blast bitscores, GC content, coverage) are transformed into criteria scores ranging from −1 to +1. Criteria scores from blast results are turned into negative values if the taxon id from the best match do not belong to a user-defined clade (such as “metazoa”). Gene scores result from the combination of criteria scores and correspond to the hyperbolic tangent of the sum of criteria scores multiplied by a ration that depends on the number of informative criteria (e.g. number of criteria for which the value is different from “0”). Scaffold scores result from the combination of gene scores and correspond to the hyperbolic tangent of the sum of gene scores multiplied by the square root of the number of genes and normalized by gene lengths. Coverage information was computed from raw ONT reads using minimap2 (parameters as follows: -ax map-ont -c -Y). Alienomics was run here under the following parameters: level_upto = metazoa; gc_filter = 26:38; evalue = 1e-01; qcovper = 0; bitscoreCutoff=150; coverage=100; ignoretaxid=104782|10195|96448. HGTs were categorized as such under the following default thresholds: genescore = 0.5; scaffoldscore = 0.5.

### Gene enrichment analyses

GO terms from functional annotation of the haploid genome (see gene annotation section above) were extracted from the 29,137 proteins predicted leading to 15,138 proteins having at least 1 associated GO term. Enrichment analyses were performed using topGO package with a fisher test and the “elim” algorithm^81^. Different genomic regions have been targeted to be compared to the rest of the genome (see Supplementary Table 2). Briefly, we compared every chromosome to the five other ones, HGTs genes to the rest of thegenes, and targeted regions on chromosome 5 with surprisingly low amount of heterozygosity in two samples (i.e. GC047403 and BXQF, see red regions in Figure 2A). Results are presented in Supplementary Table 2.

### Coverage and heterozygosity

Coverage was computed independently for the Illumina PE reads, ONT reads and PacBio reads, on 100 Kbs windows. The mapping was performed with bwa mem 0.7.17^69^ on default settings for the short-reads reads. The long reads were mapped with minimap2 2.2^59^. The coverage distribution along the 6 longest scaffolds for each read type was computed with sambamba 0.6.8^82^ (see also Supp. Fig. S9). Heterozygosity analysis was performed using GATK 4.1.0.0^83^ on Illumina PE reads for genotyping all sites (HaplotypeCaller function with -ERC GVCF option). This was done for all samples analyzed (i.e. GC047403, BXQI, BXQF, ERR321927). The resulting gvcf files were combined (CombineGVCFs function) and were then jointly genotyped (GenotypeGVCFs function)^84^. Distribution of heterozygous sites are shown on Figure 2A and on S6.

### Transposable elements analyses

Repeated elements, including transposable elements (TEs), were predicted using a combination of two complementary tools: EDTA v1.7.8^85^ and TEdenovo (part of the REPET pipeline^86, 87^. The former relies on structure-based programs allowing for the detection of even single-copy elements, while the latter relies on sequence repeatedness. The repeated elements consensus sequences they both produced were then merged to obtain 901 sequences (359 from EDTA, 542 from TEdenovo). This draft library was then automatically filtered by performing a basic annotation of the genome with TEannot from the REPET pipeline, and retrieving only consensus sequences with at least one full length copy annotated. The 652 retained consensus sequences were then used as input for the subsequent genome annotation with TEannot [REF]. This resulted in a draft annotation of 14,880 repeated elements covering 10.7% of the genome. A series of filters were then applied to these annotations using in-house script: i) conserving only retro-transposons and DNA-transposons; ii) with minimal copy length of 250 bp; iii) with minimum identity with consensus of 85%; iv) with a minimal proportion of the consensus overlapped of 33%; v) resolving overlapping annotation. These filtering steps resulted in a final annotation of 1,519 putative canonical TEs covering 3.3% of the genome. Proportions of repeated sequences and TEs are shown in Supp. Fig. S10.

### Investigating AV13 Breakpoints

The previously identified synteny breakpoints in the genome of *A. vaga* 2013 (AV13^3^) were verified by mapping the ONT reads (median size: 4,149 kb; max size: 353,147 kb) produced in this study onto the AV13 genome according to the following procedure: i) ONT reads were filtered with Porechop^88^ to discard long reads containing adapters. This discarded 1,202 out of the 1,634,477 reads; ii) Reads were mapped onto the AV13 genome using NGMLR^89^ with default parameters. This tool was selected for its accuracy when aligning long reads in a context of structural variation; iii) The scaffold of interest (i.e. scaffold1 from AV13) was aligned against the rest of the AV13 genome using Sibelia v3.0.7^90^ with the following parameters: ‘-s loose -m 10000 –gff’. iv) The haploid genome assembled here (AV19) was aligned against the AV13 genome using the same procedure as in the previous step; v) Synteny block from Sibelia were used to determine the genomic windows containing the putative breakpoints described previously^3^ (see Supplementary Table 3). These regions were manually screened using Tablet^91^ to visualize the alignment of ONT reads. We notably checked for the presence of clipped regions. Every window contained at least one clipped region (i.e. a position that is not supported by a single long read) which we reported as screenshots in S4.

### Investigating AV13 Palindromes

Palindromes previously reported in the AV13 genome were investigated in the light of our new assembly. We first *de novo* determined the location of palindromes in AV13 by filtering ONT long reads, mapping them onto AV13 genome using NGMLR^89^ with default parameters and subsequently detecting the palindromic breakpoints (PBR) using a in-house tool, huntPalindrome (available at https://github.com/jnarayan81/huntPalindrome). Each PBR location was extended by 2.5 kbp on both sides to produce PBR windows within which we checked for clipped long reads using in-house script. Additionally, we used the alignment between AV13 and AV19 genomes (as described in the previous paragraph) to show how these 20 palindromes from AV13 were assembled in AV19 (see S5). All these palindromes were collapsed into non-palindromic regions in the new AV19 genome assembly.

## Acknowledgments

We thank Alexandre Mayer for his help with scores computation and transformation within Alienomics, Mathilde Colinet for her help with FISH experiments as well as Nathalie Verbruggen for kindly providing *Arabidopsis* plantlets for genome size estimation.

## 1 Supplementary Materials

**Supplementary Figure S1.**
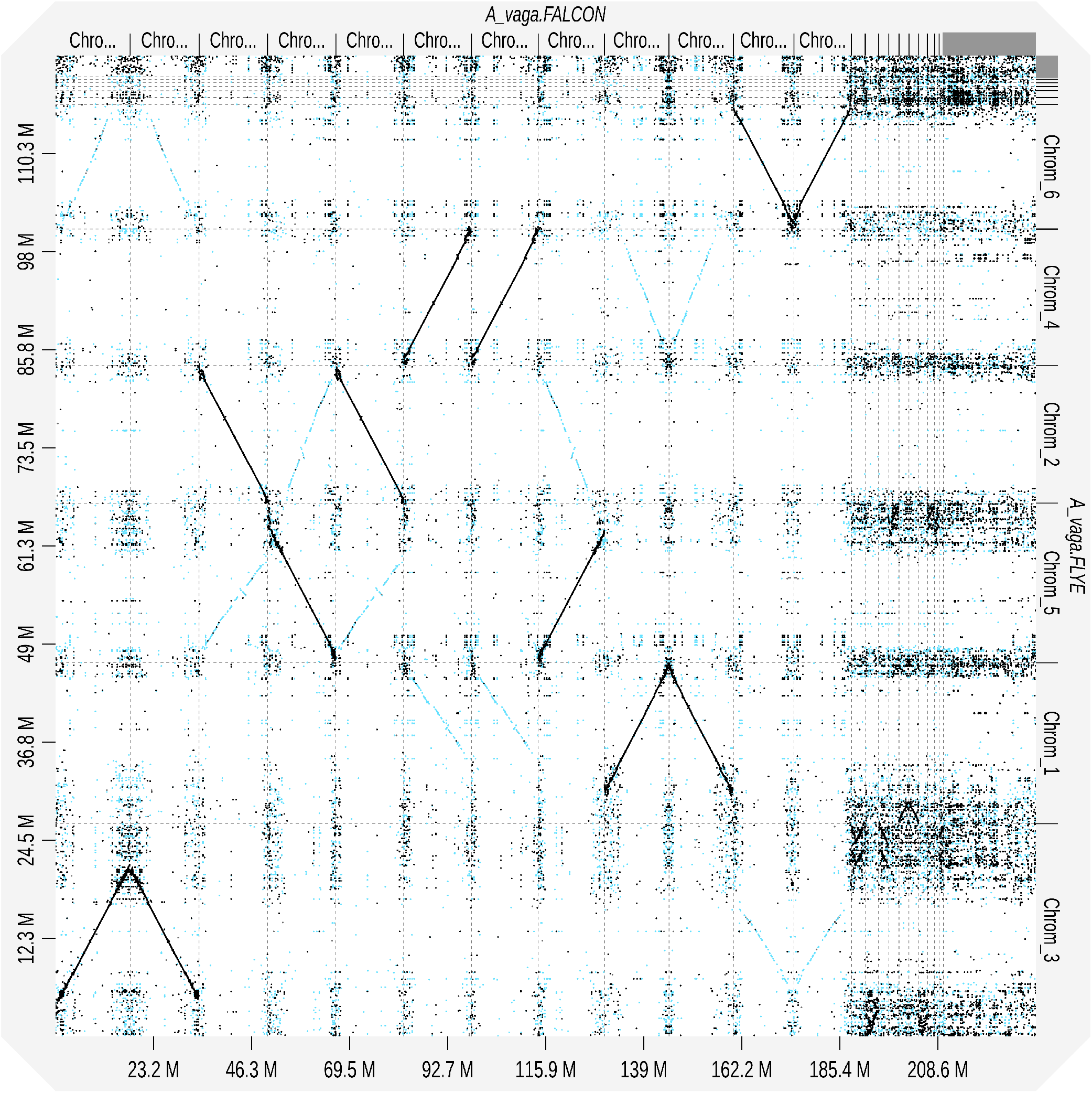
Haploid versus diploid genome dotplot. Pairwise alignment of the haploid assembly (Flye) against the diploid assembly (Falcon), vizualized using D-GENIES.

**Supplementary Figure S2.**
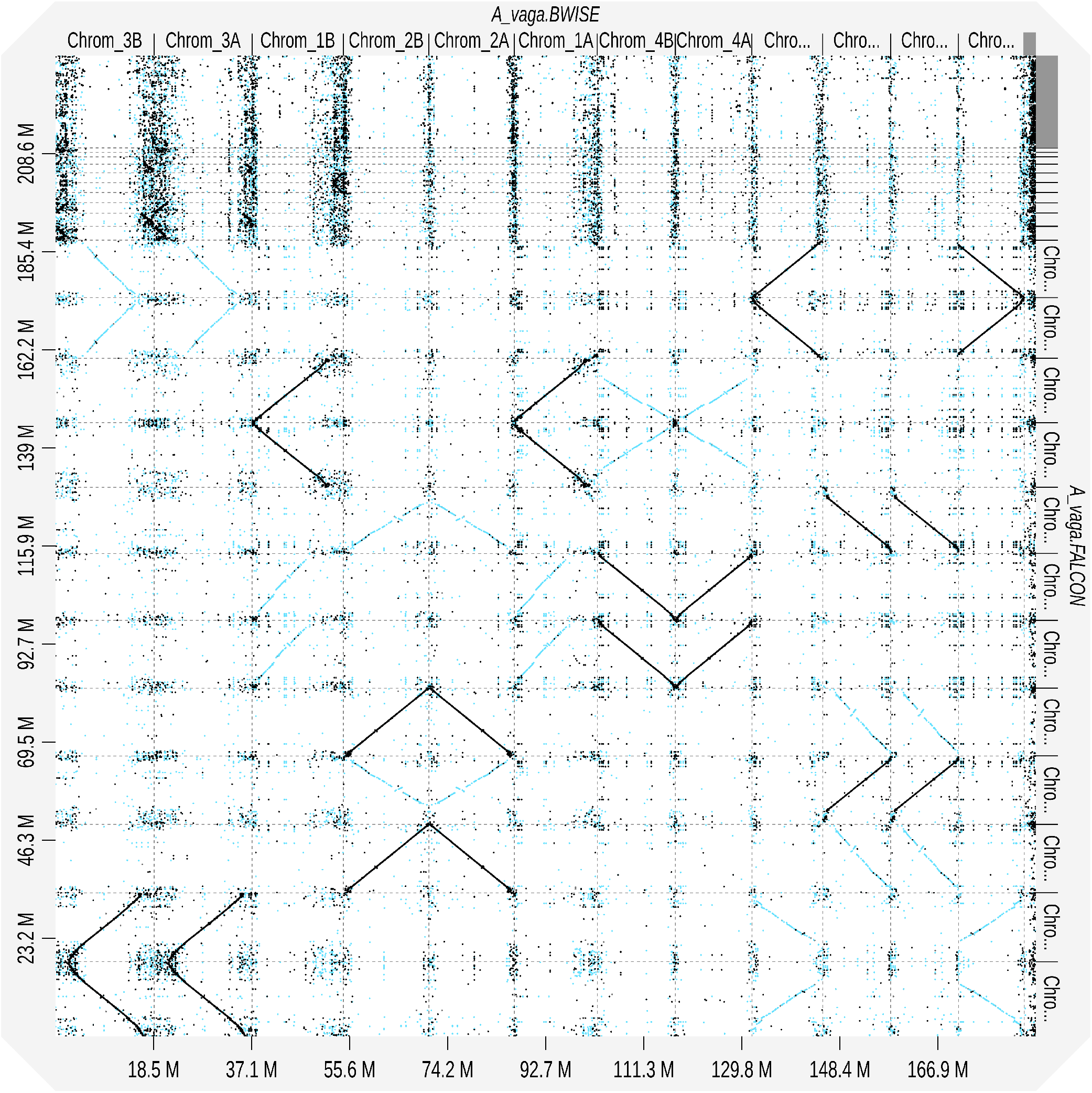
diploid versus phased genome dotplot. Pairwise alignment of the diploid assembly (Falcon) against the free assembly (BWISE), vizualized using D-GENIES.

**Supplementary Figure S3.**
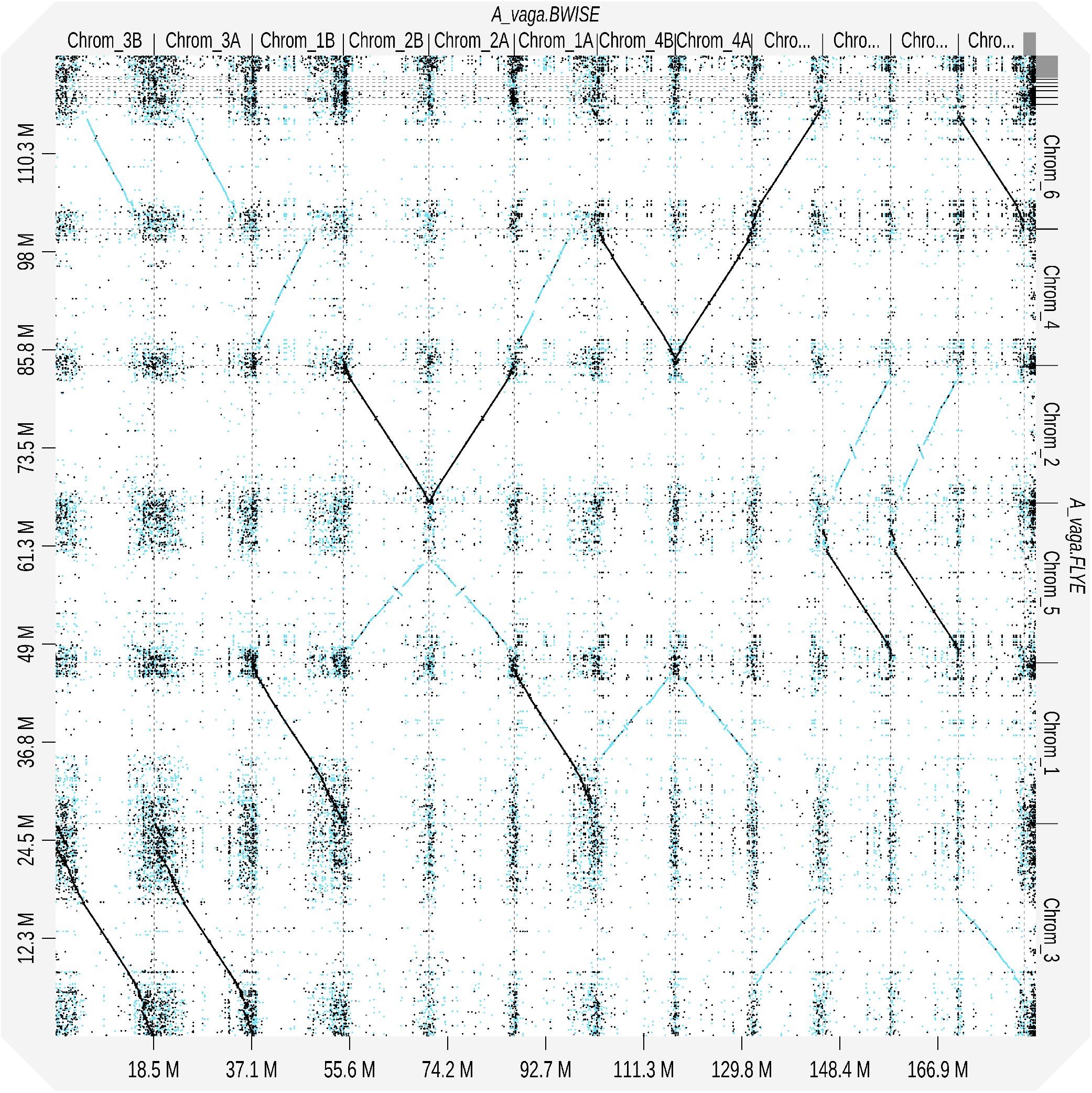
Haploid versus phased genome dotplot. Pairwise alignment of the haploid assembly (Flye) against the free assembly (BWISE), vizualized using D-GENIES.

**Supplementary Figure S4.**
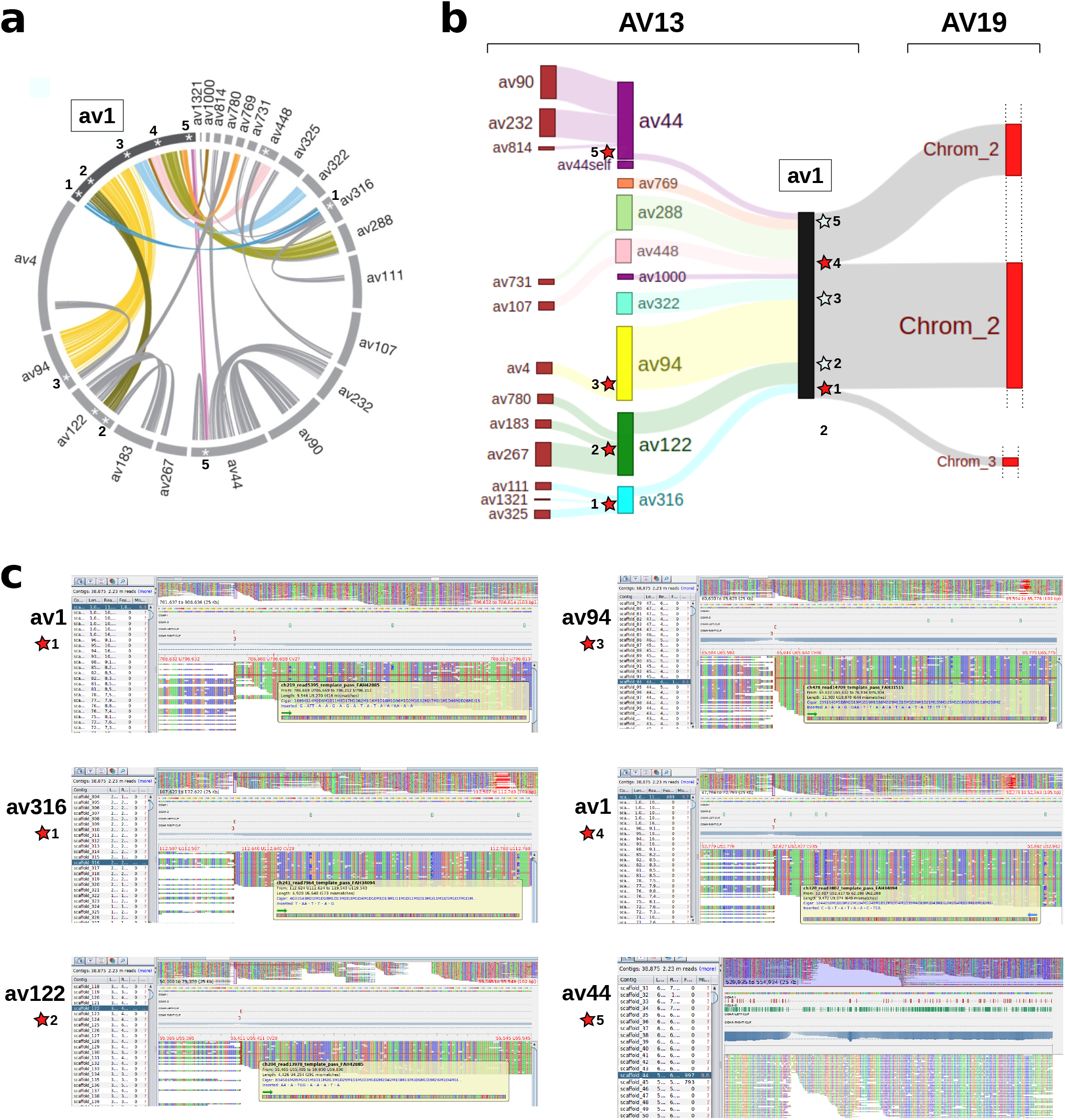
Invalidating AV13 breakpoints. Schematic of the structural differences identified between AV13 scaffold1 and AV19. a) Schematic view of AV13 genome synteny depicting the 5 five putative colinear breakpoints on the scaffold av1 (adapted from Flot 2013). b) Schematic view of synteny alignment between AV13 and AV19, depicting the 5 putative colinear breakpoints of scaffold av1. Red stars indicate genomic region that are not supported by long reads, while white stars indicate regions supported by long reads. c) Screenshots of long read alignments from Tablet depicting clipped reads that indicate a problem in the AV13 assembly.

**Supplementary Figure S5.**
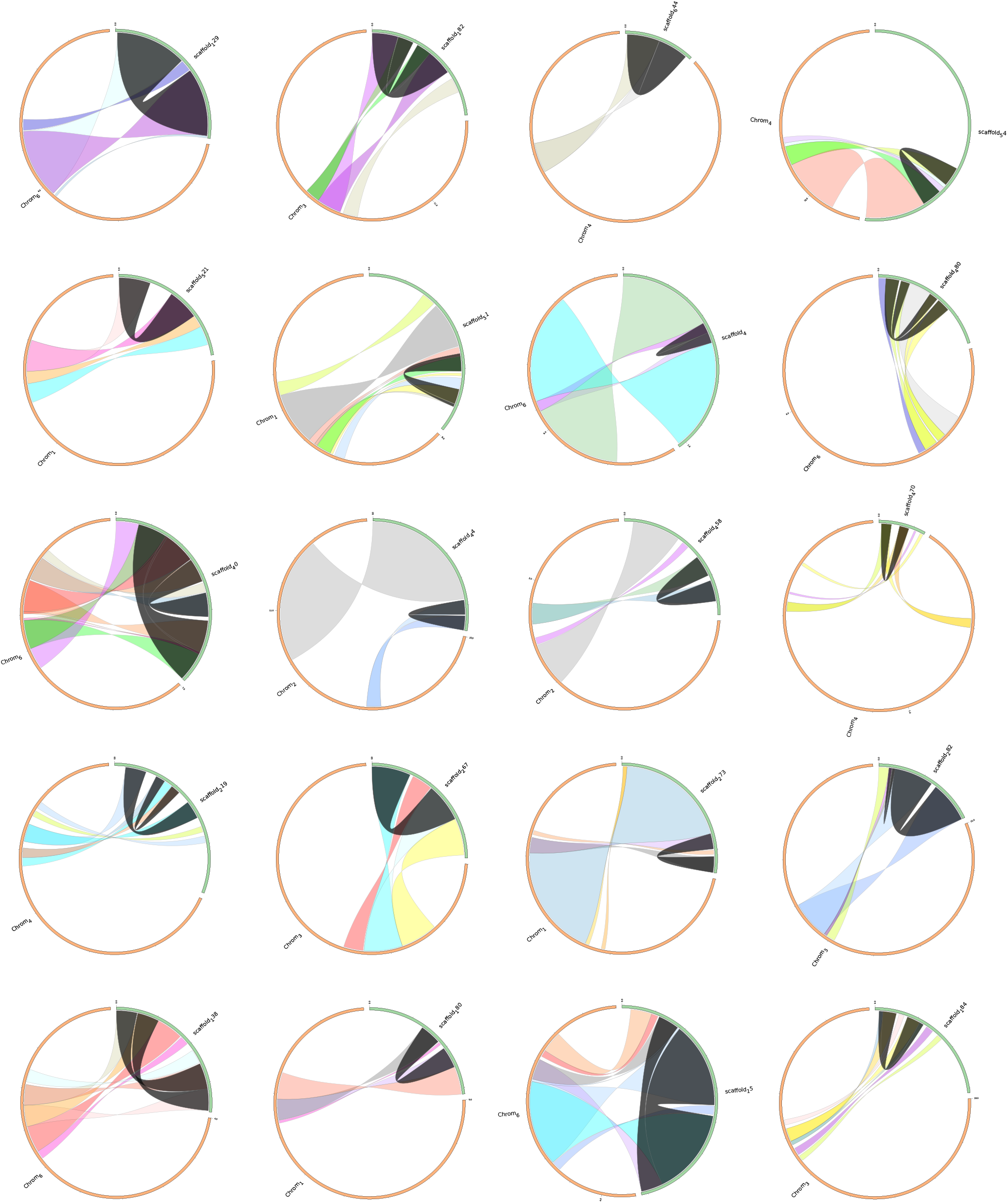
Invalidating AV13 palindromes. Alignment of the AV13 genome assembly^3^ against our new AV19 genome assembly shows the total absence of previously reported palindromes. Green bars represent scaffolds from 2013 assembly and orange bars represents chromosomes assembled in the present study. Palindromic regions in 2013 assembly are shown in dark grey.

**Supplementary Figure S6.**
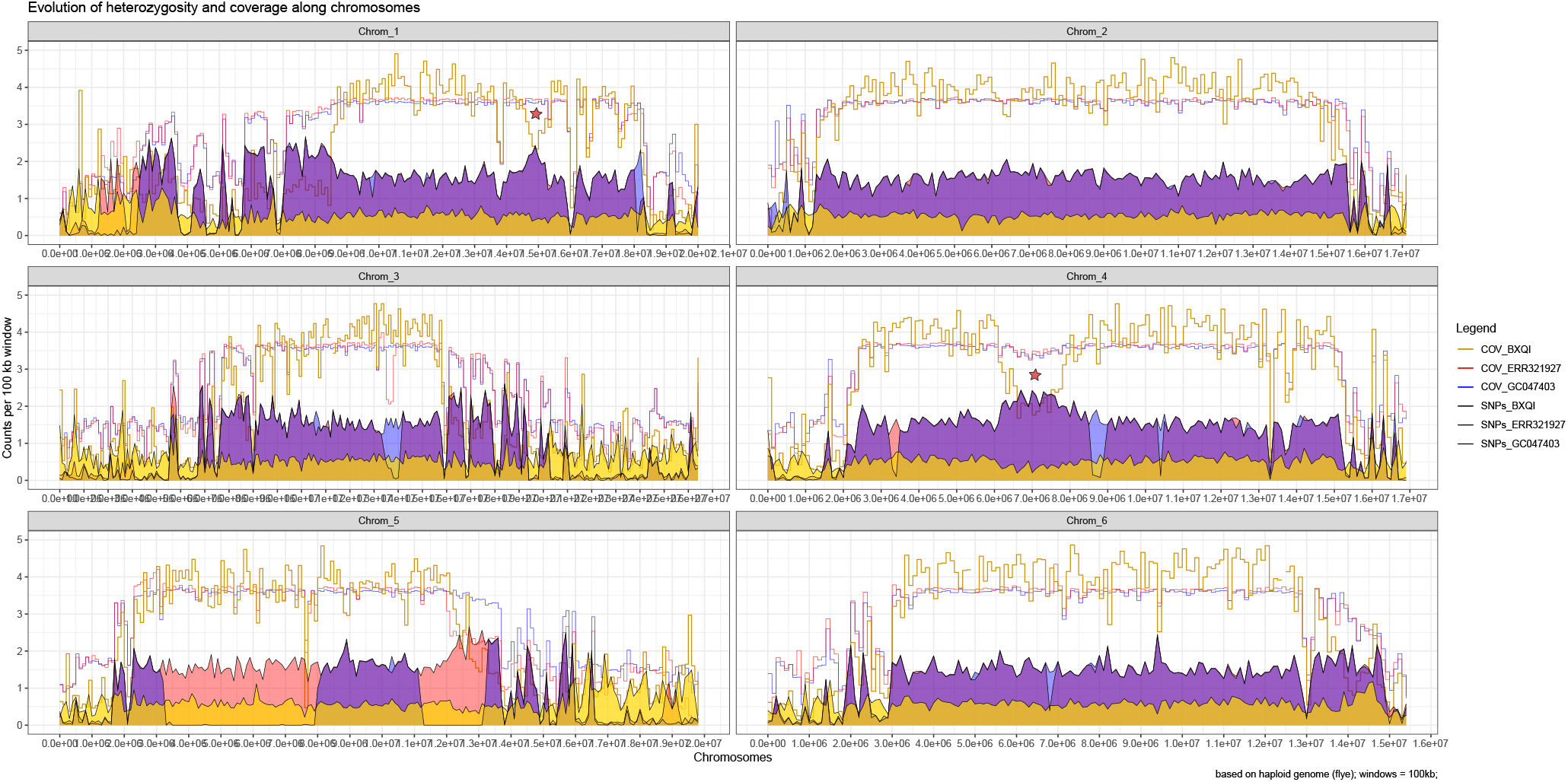
Coverage and heterozygosity in the AV19 genome assembly. Heterozygosity changes along the six chromosomes including a wild *A. vaga* sample (i.e. BXQI).

**Supplementary Figure S7.**
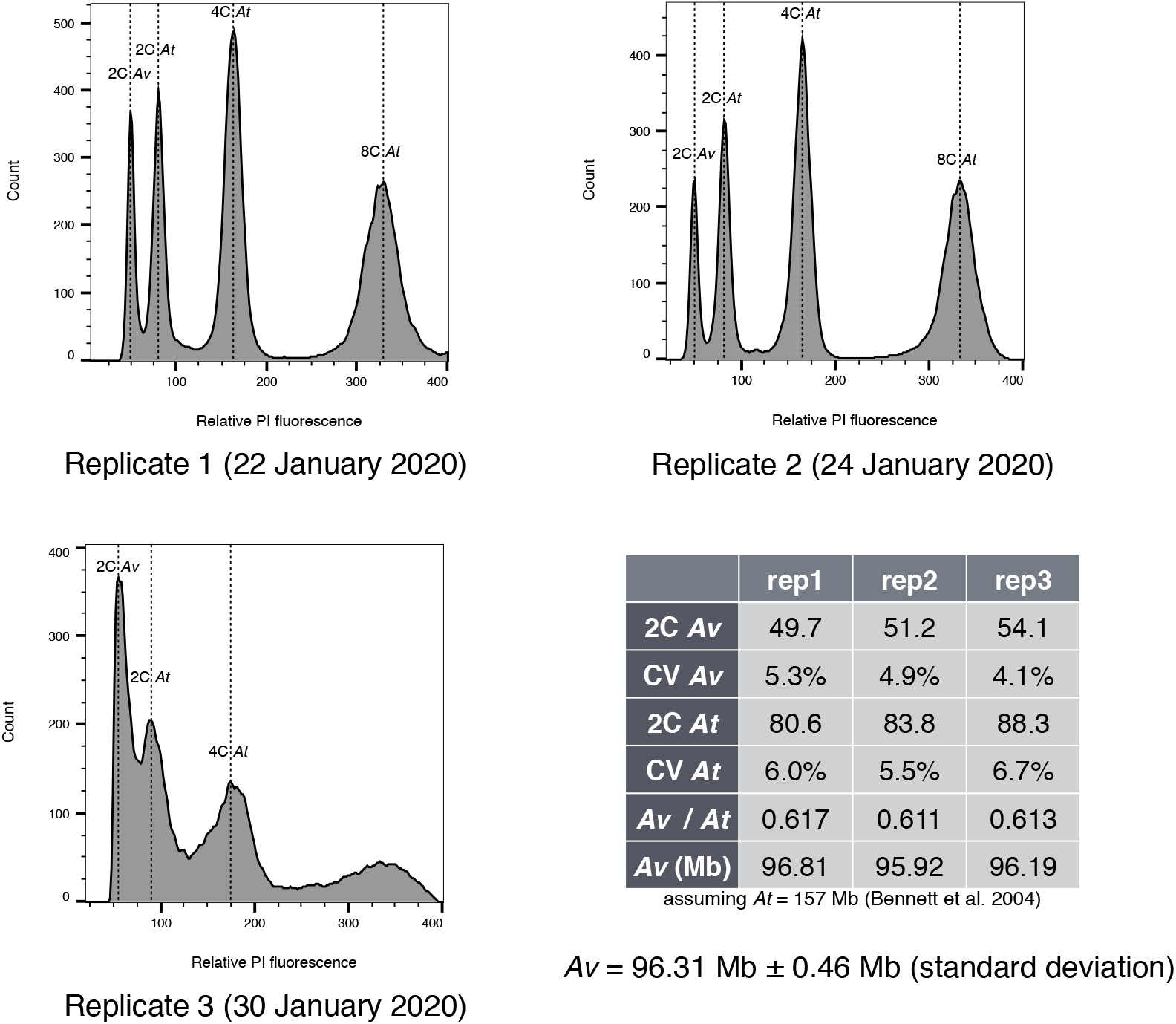
Genome size estimation. Flow cytometry measurement of the genome size of *Adineta vaga* (*Av*) by comparison to *Arabidopsis thaliana* cultivar Colombia (*At*).

**Supplementary Figure S8.**
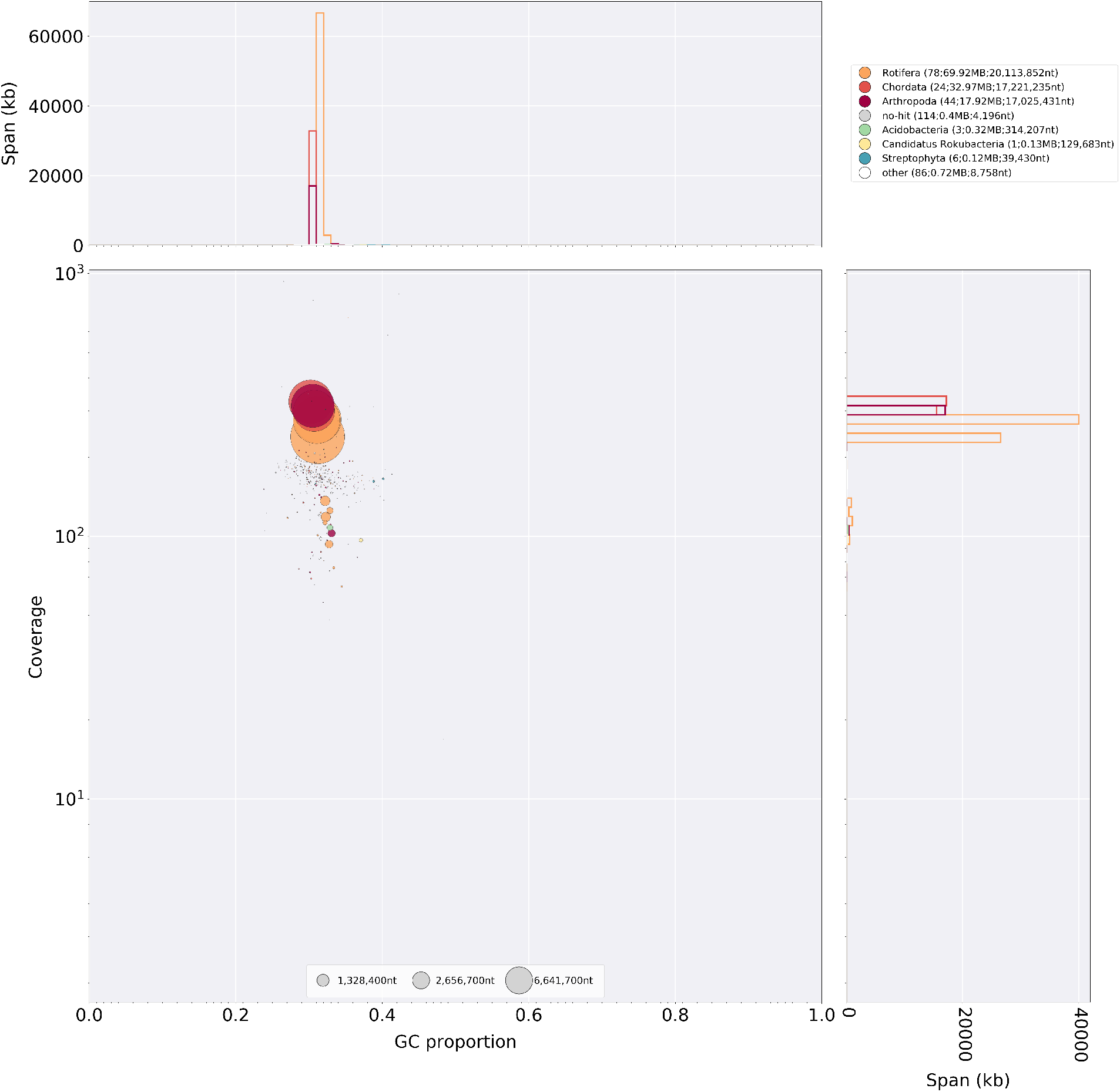
Contamination check with blobtools. Blobplot of the haploid assembly (Flye). Circles represent scaffolds, the coordinates are determined by the average coverage and GC content, the diameter represents scaffold size and the color corresponds to taxonomic assignment.

**Supplementary Figure S9.**
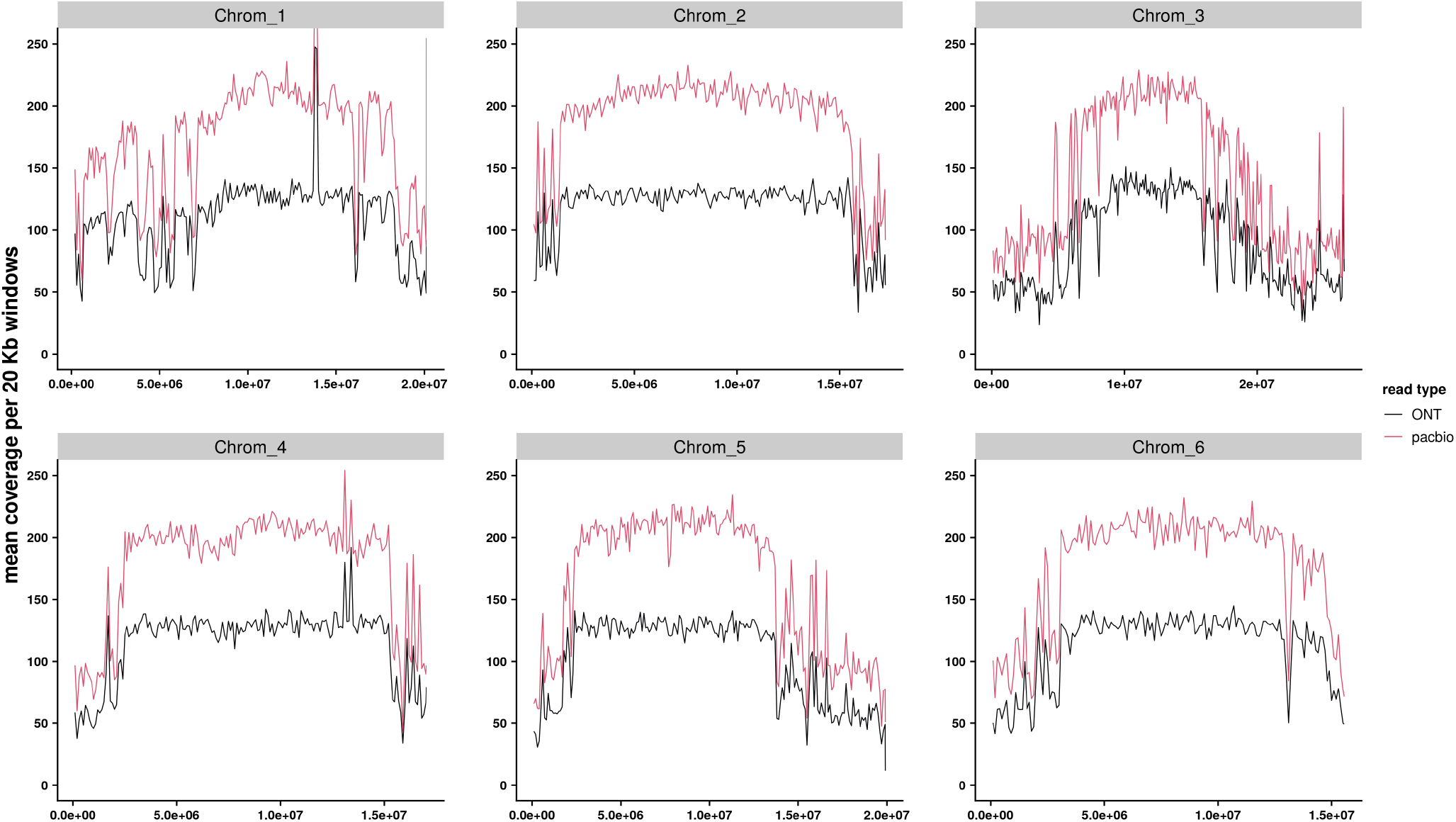
Long reads coverage. Coverage of long reads along the 6 chromosomes of AV19 genome. Red line: PacBio reads. Black line: ONT reads.

**Supplementary Figure S10.**
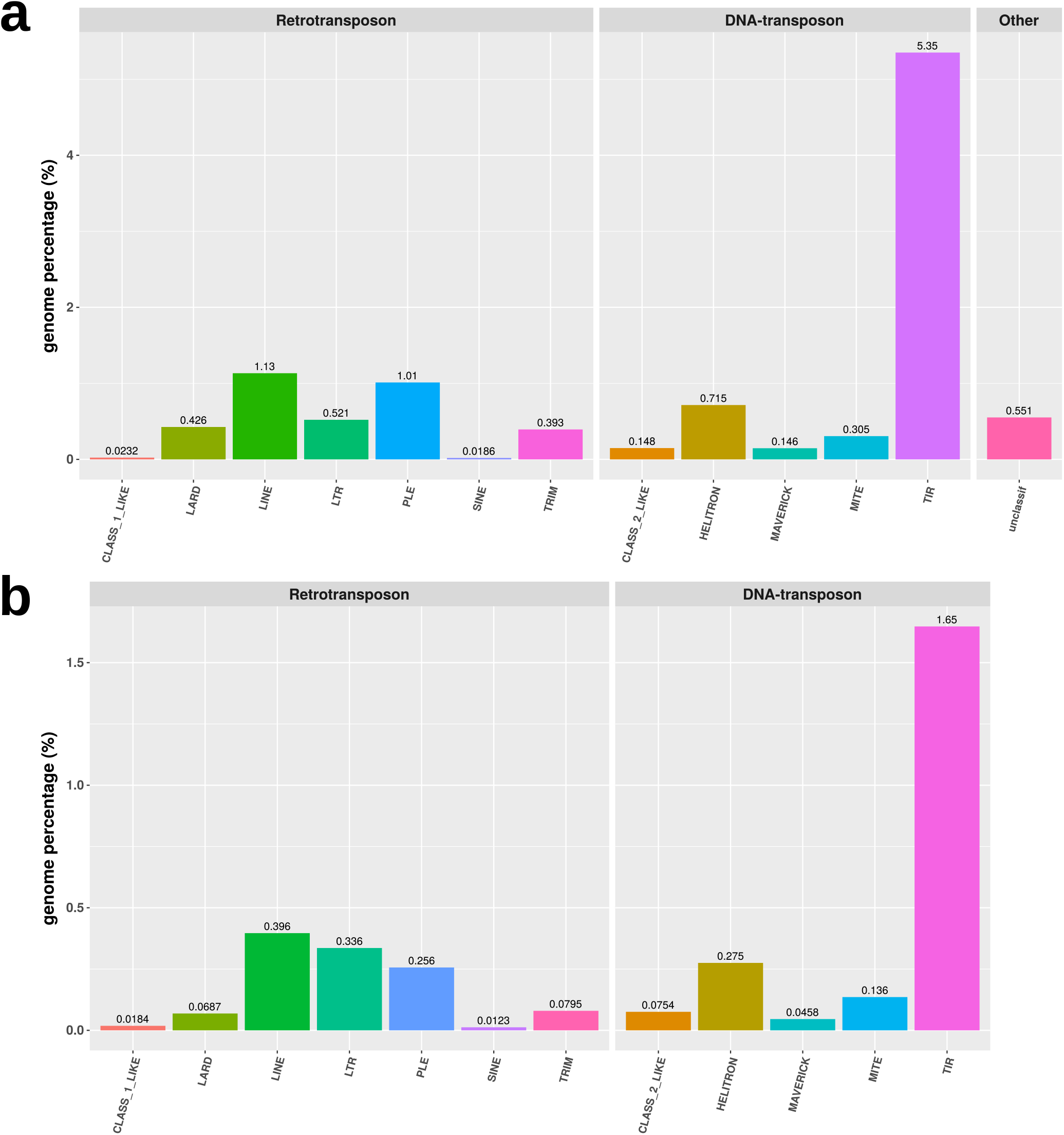
Transposable elements analyses. Proportion of the genome covered by each TE class for: a) draft annotation including all repeated elements; b) filtered annotation, including only putative canonical TEs.

